# Cortical Circuit Mechanisms of Multimodal Temporal Pattern Discrimination

**DOI:** 10.1101/2022.08.31.506133

**Authors:** Sam Post, William Mol, Omar Abu-Wishah, Shazia Ali, Noorhan Rahmatullah, Anubhuti Goel

## Abstract

Discriminating between temporal features in sensory stimuli is critical to complex behavior and decision making. However, how sensory cortical circuit mechanisms contribute to discrimination between subsecond temporal components in sensory events is unclear. To elucidate the mechanistic underpinnings of timing in primary visual cortex (V1), we recorded from V1 using 2-photon calcium imaging in awake-behaving mice performing a go/no-go discrimination timing task, which was composed of patterns of subsecond audio-visual stimuli. In both conditions, activity during the early stimulus period was temporally coordinated with the preferred stimulus. However, while network activity increased in the preferred condition, network activity was increasingly suppressed in the nonpreferred condition over the stimulus period. Our results demonstrate that discrimination between subsecond intervals that are contained in rhythmic patterns can be accomplished by local networks and suggest the contribution of neural resonance as a mechanism.

## Introduction

A key aspect of sensory discrimination in learning and memory and generating complex behavior is extracting temporal features from external stimuli. Particularly in the range of seconds and subseconds, the accurate encoding of durations and sequences of durations influences a variety of sensory and motor abilities. For example, one may need to keep a beat and synchronize tempo when in a band; a prey may need to jump out of the way of a predator at just the right moment; the timing of a yellow light must be predicted in order to decide whether to slow down or to go through it; and meaning in spoken language derive s from sequences of syllables that are highly temporally structured. Based on psychophysical and pharmacological data, it is most likely that there are multiple neural mechanisms that code for temporal structure of sensory events since they are timed over a broad range of timescales, ranging from microseconds to days (Mauk and Buonomano, 2004; Buhusi and Meck, 2005; Paton and Buonomano, 2018). However a growing body of literature suggests that time intervals in the sub-second and second range are encoded in the emergent changing patterns or neural dynamics across many brain areas, including sensory cortex (Pastalkova et al., 2008; Carnevale et al., 2015; Gouvea et al., 2015; Namboodiri et al., 2015; Goel and Buonomano, 2016; Soares et al., 2016; Bakhurin et al., 2017; Emmons et al., 2017; Heys and Dombeck, 2018; Tsao et al., 2018; Zhou et al., 2020).

Temporal structure in sensory stimuli is often organized as rhythms which can be simple isochronous sequences in which all the intervals are identical, or which can be a complex combination of time intervals. However, the neural dynamical regimes in the sensory cortex that contribute to processing and learning rhythmic patterns in external stimuli remain largely unclear: Specifically, do emergent neural dynamics in primary visual cortex (V1) contribute to learning the temporal structure of rhythmic patterns? One prevailing idea is that neural oscillations allow communication between sensory and motor cortical areas, thus producing temporal predictions and entrainment to rhythms (Merchant et al., 2015). Our goal is to understand how visual cortical dynamics adapt to the temporal structure of a multimodal rhythm in a goal directed task.

We investigate rhythmic patterns in V1 based upon a large body of experimental evidence that shows that primary visual cortical circuits show robust plasticity to spatiotemporal features in stimuli and predict temporal associations (Chubykin et al., 2013; Gavornik and Bear, 2014b, a; Fiser et al., 2016; Garrett et al., 2020). Further, visual cortical plasticity is influenced by functional input from other brain areas such as hippocampus (Finnie et al., 2021) and auditory cortex (McIntosh et al., 1998; Zangenehpour and Zatorre, 2010; Deneux et al., 2019; Garner and Keller, 2022) and audio-visual stimuli evoke multimodal plasticity in primary visual cortex (V1) (Morrell, 1972; Petro et al., 2017).

We bring our goal to an *in vivo* setting by implementing a novel audio-visual timing task, temporal pattern sensory discrimination (TPSD), in awake behaving mice using 2-photon calcium imaging in V1, layer 2/3 (L-2/3). In the TPSD task mice learn to discriminate between two temporal patterns. Our paradigm builds upon previous work in temporal pattern discrimination which suggests that multisensory stimuli enhance discriminability of sequences (Raposo et al., 2012; Barakat, Sietz, and Sham, 2015). This project will be highly impactful for the following reasons. 1) Sensory timing tasks in animal models like mice allow for more granular recording methods and a broader and more nuanced toolkit of experimental manipulations. 2) To delineate the emergent dynamical mechanisms that encode information about rhythmic durations in a goal driven task. Both reasons will improve our fundamental understanding of how information about temporal patterns is extracted from sensory stimuli and learned, and will also provide insights into timing deficits in neurological disorders.

## Results

### Mice learn to perform a multimodal temporal pattern sensory discrimination task

To examine temporal pattern learning we have designed a novel go/no-go, Temporal Pattern Sensory Discrimination (TPSD) task (see methods). We test our paradigm in mice as they are a robust animal model for temporally and spatially fine recording methods and cell-type specific tagging and manipulation. Awake, head-restrained young adult mice (2-3 months) are habituated to run on a polystyrene ball treadmill while they perform the TPSD paradigm. Water-deprived mice are presented with two audio-visual temporal patterns (preferred and non-preferred) as shown in the schematic in **Fig. 1B**. Each pattern consists of 4 audio-visual (AV) stimuli, where each AV stimulus lasts either 0.2s or 0.9s and is separated by a 0.2s gray screen. The visual stimulus consists of a drifting sinusoidal 90° grating, and the auditory stimulus consists of a 5kHz tone. Both auditory and visual stimuli are presented concurrently; therefore, the stimuli are audio-visual. The temporal pattern with 0.2s AV stimuli is associated with a water reward (preferred pattern) and the temporal pattern with 0.9s AV stimuli is not (non-preferred pattern) (**Fig. 1B**). We quantified performance of mice using a discriminability index (*d’*) in which 2 was set as a learning threshold (**Fig. 1C**). Mice learn to preferentially lick to the preferred pattern and withhold licking for the non-preferred pattern (11 ± 2.58 sessions to learn; one-way ANOVA, *F*_1,13_ = 8.91, p = 6 × 10^−10^).

**Figure 1.**
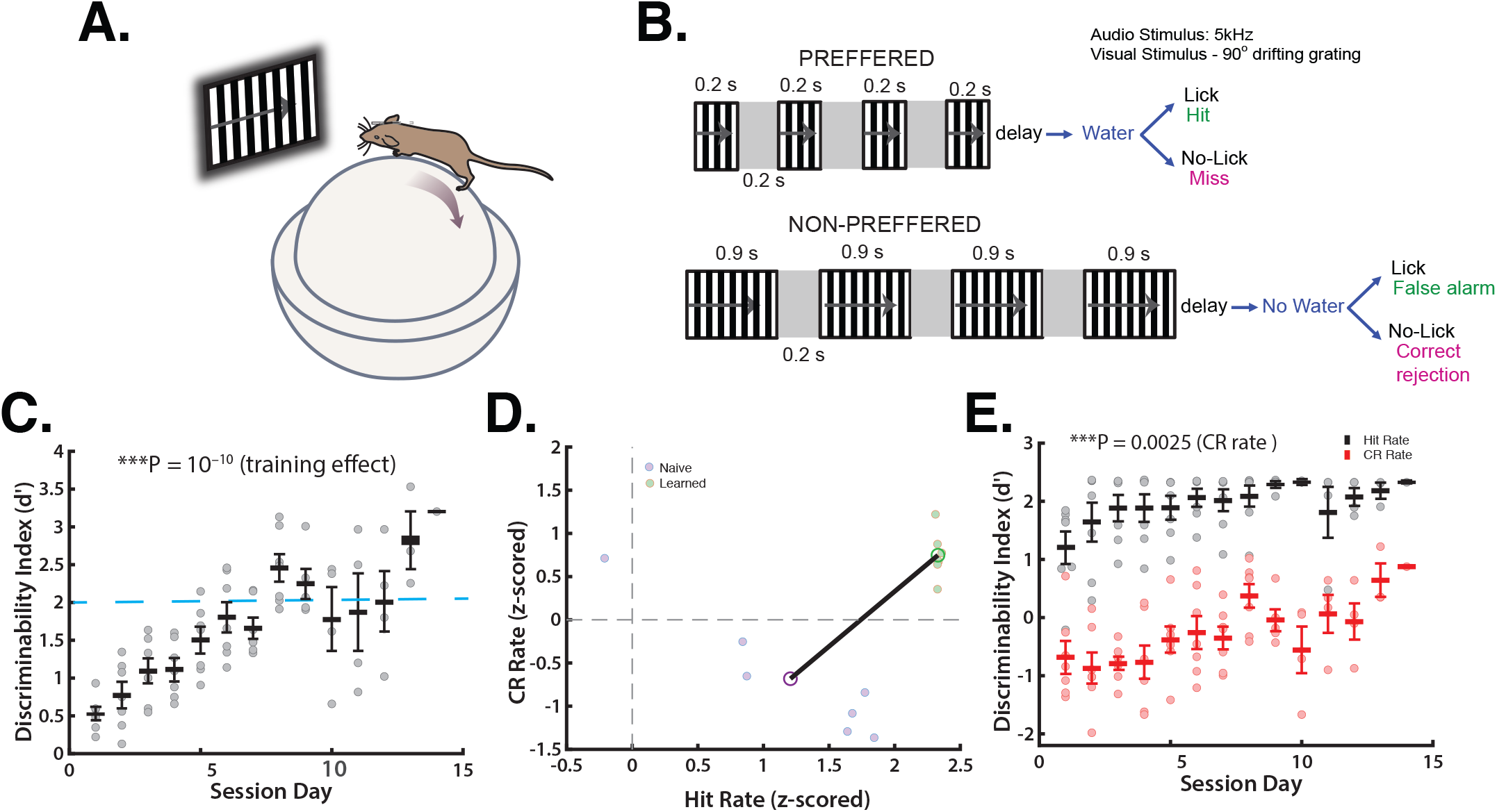
Mice learn to perform a multimodal temporal pattern sensory discrimination task. **A**. Schematic of behavior task in a head fixed mouse on a polystyrene ball. **B**. Timeline of an individual trial of the go/no-go Temporal Pattern Sensory Discrimination task. **C**. Discriminability index (d’) shows mice learn to discriminate temporal patterns (one-way ANOVA, *F*_1,13_ = 8.91, p = 6 × 10^−10^). The dashed line at d′ = 2 indicates the expert performance threshold. **D**. Hit rates and Correct Rejection (CR) rates both change with learning (Hit rate: Wilcoxon rank sum test; n = 7, *p* = 5.83×10^−4^. CR rate: two-sided t-test; n = 7, *t*(12) = 4.9033, *p* = 3.64×10^−4^). **E**. Increased performance is associated with a significantly higher proportion of CR responses. (one-way ANOVA, *F*_1,13_ = 2.89, p = .0025) rather than HR rate (Kruskal-Wallis test, *H*(13) = 16.28, *p* = .23).

To confirm that learning is not simply a biased feature of the differential preferred-nonpreferred ratio (7:3) and to examine the decision strategy of mice between learned and naive days we compared Hit rates (Hr) with Correct Rejection rates (CRr) (**Fig. 1D**). Mice improved performance primarily by improving their CRr in which their CRr changed from negative to positive (two tailed, paired sample Student’s *t*-test, *t*(12) = 4.9033, p = 3.64×10^−4^). Although significant, we find that mice’s Hr remains relatively unchanged across sessions (Wilcoxon rank sum test, p = 5.83×10^−4^) but that their ability to inhibit licking increases across sessions, indicated by an increased CRr. We additionally examined changes in Hr and CRr across sessions and found a significant difference in CRr (one-way ANOVA, *F*_1,13_ = 2.89, p = .0025) but not Hr (Kruskal-Wallis test, *H*(13) = 16.28, *p* = .23), indicative of changes in CRr driving successful performance (**Fig. 1E**).

Licking profiles that accompany learning were more refined in expert mice (**Fig. 2A**). We next quantified probabilities and cumulative distributions of mice’s licking based on session day (naive vs. learned) and stimulus type (preferred vs. nonpreferred) as a function of time. We find that on learned days licking to the preferred stimulus is enhanced, while licking to nonpreferred stimuli dramatically decreases prior to the water reward, indicative of a learned response (**Fig. 2B,C**) (**Fig 2C:** two-sided Kolmogorov-Smirnov tests with Bonferroni Correction: new Alpha = .0125. Pref naive vs. NP naive: *D*(101225) = .0099, *p* = .0336. Pref learned vs NP learned: *D*(86883) = .1376, *p* = 3.17×10^−276^. Pref naive vs. pref learned: *D*(135670) = .1027, *p* = 7.57×10^−311^. NP naive vs. NP learned: *D*(52438) = .0613, *p* = 1.87×10^−2^). The most robust change in licking occurred in the nonpreferred stimulus in which mice peak in their lick probabilities prior to onset of the water reward in learned days (**Fig. 2B,D**).

**Figure 2.**
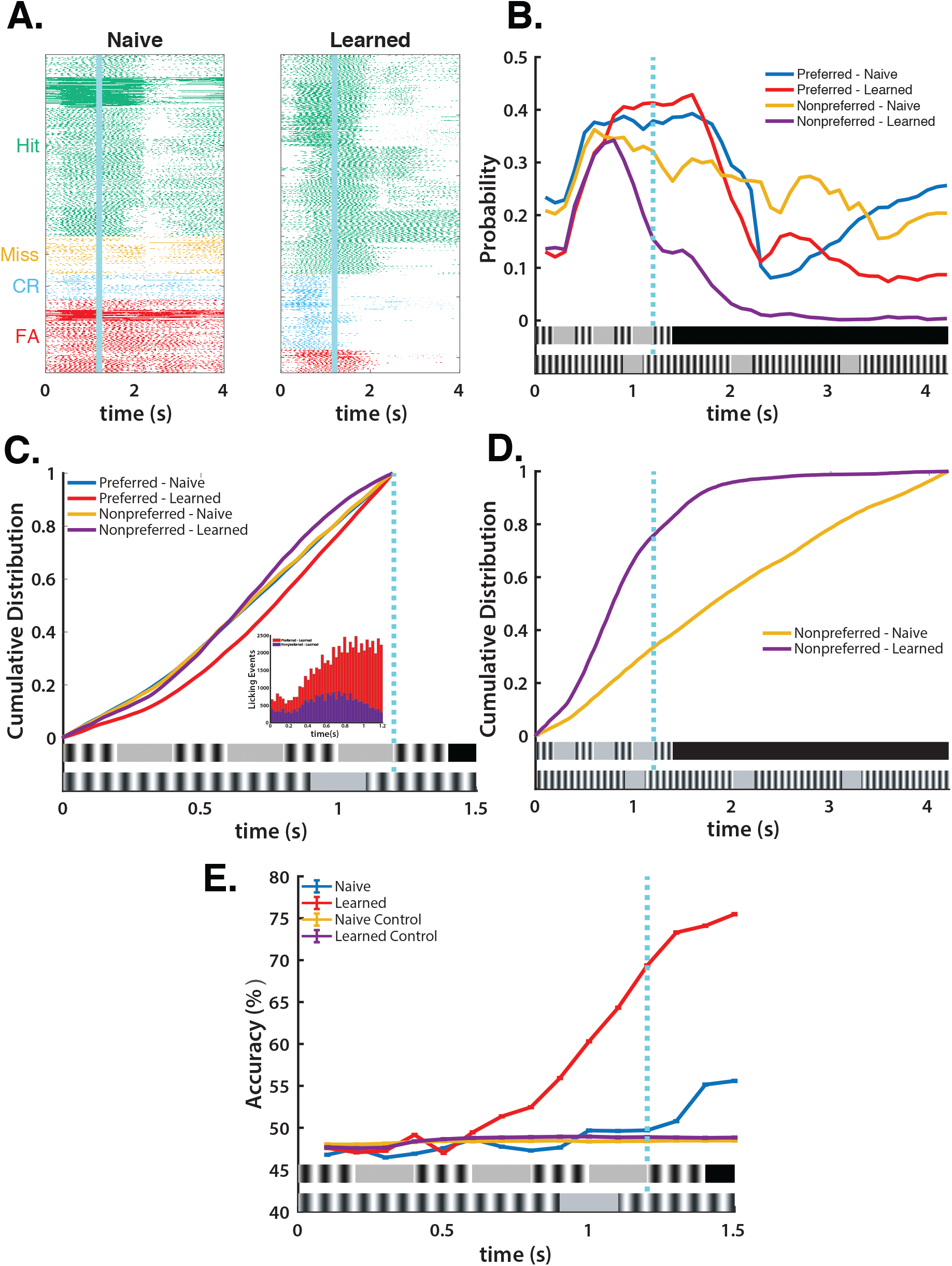
Licking Profiles in learned mice predict stimulus type. **A**. Raster of plots of licking of all mice (n=7) for each’s best 150 trials in the naive and learned days. **B**. Probabilities of licking based on session day and stimulus type. Nonpreferred stimulus licking decreases prior to the water reward (dashed blue line) at 1.2 s on the learned day. **C**. Cumulative Distribution of licking from stimulus onset to water reward at 1.2 s (two-sided Kolmogorov-Smirnov tests with Bonferroni Correction: new Alpha = .0125. Pref naive vs. NP naive: *D*(101225) = .0099, *p* = .0336. Pref learned vs NP learned: *D*(86883) = .1376, *p* = 3.17×10^−276^. Pref naive vs. pref learned: *D*(135670) = .1027, *p* = 7.57×10^−311^. NP naive vs. NP learned: *D*(52438) = .0613, *p* = 1.87×10^−2^). Inset of histogram showing raw licking histogram in learned preferred and nonpreferred stimuli for all mice (0 to 1.2 s). **D**. Cumulative Distribution of licking from stimulus onset to end of nonpreferred stimulus (0 to 4.2 s) (two-sided Kolmogorov-Smirnov test; NP naive vs. NP learned: *D* (118971) = .4516, *p* → 0). **E**. Support Vector Machine (SVM) increasingly predicts stimulus from licking in the learned but not naïve days prior to the water reward.

### Licking Profiles in learned mice predict stimulus type

To establish more causally that licking is both 1) a viable measure of performance; and 2) demonstrates differential learning between mice, we developed a bootstrapped Support Vector Machine (SVM), a type of binary classifier. SVMs in effect generate coordinates for infinite dimensions from low dimensional coordinates; this allows them to exaggerate the Euclidean space between the original low dimensional points such that low dimensional points of high similarity retain high similarity and low dimensional points of low similarity increase their dissimilarity. We run our SVM 10,000 times within 0.1 s time bins using licking within a trial and the stimulus of that trial as the outcome (see Methods). This allows us to generate a distribution of correctly predicted outcomes per time bin per mouse, which are then compared to a stimulus-shuffled control. We find that there is little predictability beyond chance in naive days in mice with somewhat greater predictability following the water reward, likely attributable to increased licking at their chance encounter with the water reward (**Fig. 2E**). On learned days, licking becomes predictive beyond chance at 0.7 s and then accelerates beginning at 0.8 s. This suggests that mice are relying on stimulus information to make a decision, rather than the presence or absence of the water reward. The SVMs’ high performance prior to the water reward in learned sessions establishes that mice indeed learn to discriminate temporal patterns. In addition, it identifies the decision period at approximately 0.7-0.8 s, which is the period immediately following the conclusion of the second stimulus in the preferred stimulus.

### Pyramidal cell dynamics in primary visual cortex accompany temporal pattern learning

In our published study using sensory cortex tissue (Goel and Buonomano, 2016) we found that information about stimulus duration is encoded in a change in pyramidal cell activity wherein the neural activity is refined to represent the learned interval. Studies have shown that auditory inputs strongly influence neural responses in primary visual cortex (McIntosh et al., 1998; Zangenehpour and Zatorre, 2010; Deneux et al., 2019; Garner and Keller, 2022) and audio-visual stimuli evoke multimodal plasticity in V1 (Morrell, 1972; Petro et al., 2017). Therefore, to examine the pyramidal cell dynamics that are associated with TPSD, we performed two photon calcium imaging in V1 to provide a real time assay of neural activity during TPSD. We recorded from V1, L-2/3 using 2-photon calcium imaging using jGCaMP7f (half-rise time 27±2 ms) with a synapsin promotor via an AAV vector (Dana et al., 2019) (**Fig. 3-D**). This indicator has been utilized by numerous published studies and is routinely used by researchers performing calcium imaging during behavior, due to its enhanced signal to noise ratio and fast rise time kinetics.

**Figure 3.**
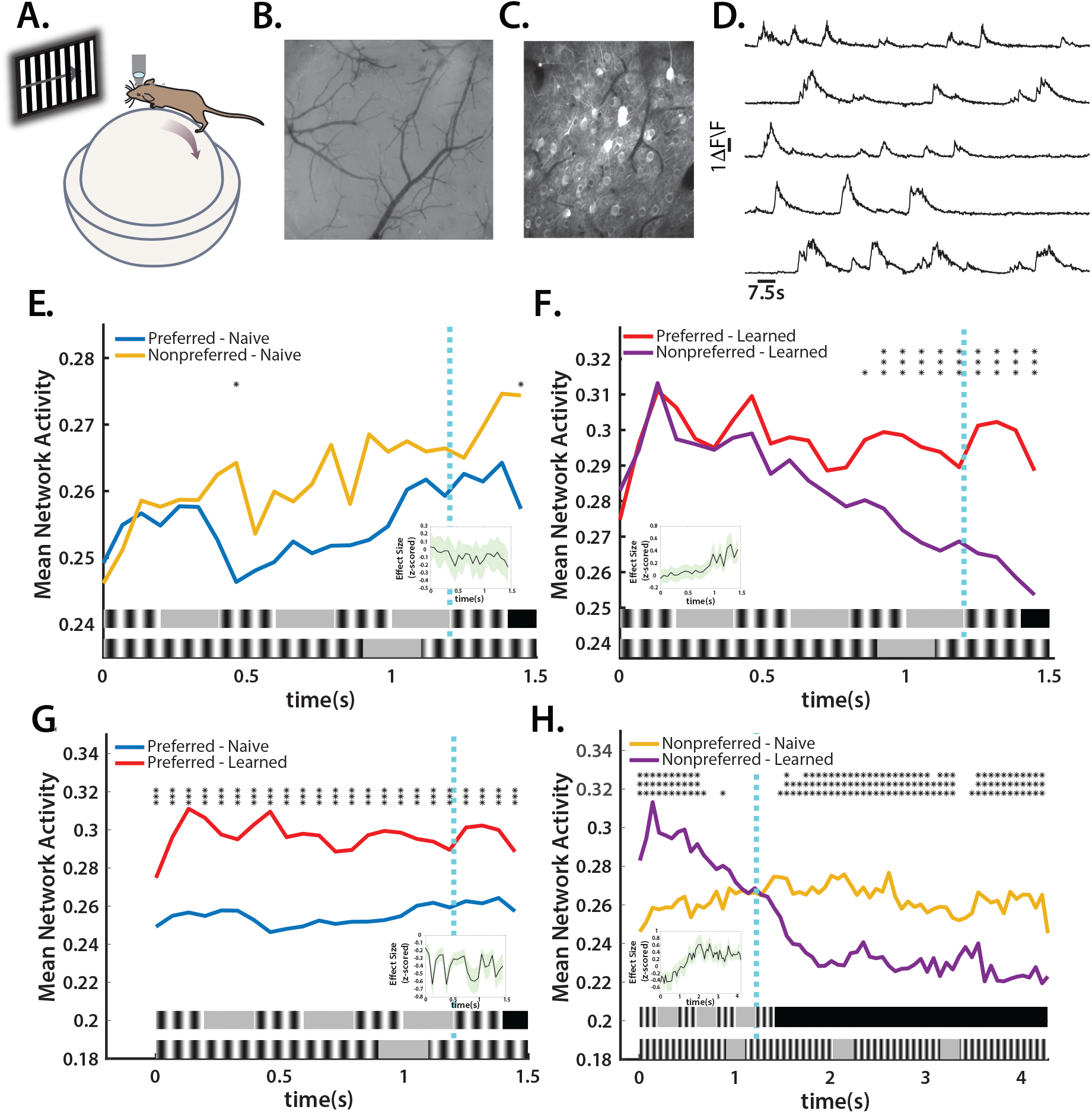
Pyramidal cell dynamics in primary visual cortex accompany temporal pattern learning. **A**. Schematic of calcium imaging during the behavior task in a head fixed mouse. **B**. Craniotomy over V1. **B**. Representative frame of video taken from binocular V1 during behavior with imaging. **C**. Representative field of view for in vivo two-photon calcium imaging experiment in V1.**D**. Example traces of changes in jGCaMP7f fluorescence intensity (Δ F/F) for 5 representative neurons in V1. **E**. Mean network activity averaged over trials in naive days does not show temporal coordination or differentiation (23 .067 s bins were compared using either a two-tailed t-test or a Wilcoxon rank sum test following a Lilliefors test to test normality per time bin, with a Bonferroni Correction: new Alpha value = .0022; inset shows effect sizes with confidence intervals at each time bin using either Cohen’s D or Cliff’s delta following Lilliefors test to determine normality.) **F**. Mean network activity averaged over trials in learned days shows temporal coordination and differentiation (23 .067 s bins were compared using either a two-tailed t-test or a Wilcoxon rank sum test following a Lilliefors test to test normality per time bin, with a Bonferroni Correction: new Alpha value = .0022; inset shows effect sizes with confidence intervals at each time bin using either Cohen’s D or Cliff’s delta following Lilliefors test to determine normality). **G**. Mean network activity averaged over trials across preferred stimulus period shows elevated activity in learned days (23 .067 s bins were compared using either a two-tailed t-test or a Wilcoxon rank sum test following a Lilliefors test to test normality per time bin, with a Bonferroni Correction: new Alpha value = .0022; inset shows effect sizes with confidence intervals at each time bin using either Cohen’s D or Cliff’s delta following Lilliefors test to determine normality.) **H**. Mean network activity averaged over trials across nonpreferred stimulus period shows initial elevated activity in learned days followed by decreasing activity following the end of the preferred stimulus period (65 .067 s bins were compared using either a two-tailed t-test or a Wilcoxon rank sum test following a Lilliefors test to test normality per time bin, with a Bonferroni Correction: new Alpha value = 7.69×10^−4^; inset shows effect sizes with confidence intervals at each time bin using either Cohen’s D or Cliff’s delta following Lilliefors test to determine normality.)

Our hypothesis is that during the course of training, learning will be mediated by an enhancement of pyramidal cell activity in V1. Specifically, similar to our *in vitro* studies (Johnson et al., 2010; Goel and Buonomano, 2016), we expect to see distinct patterns or distributions of activity emerge that encode information about each temporal pattern. We imaged from mice (n = 4) performing the TPSD task. We find that in mice during the TPSD task, there are changes in mean activity of the network in a time-dependent manner (**Fig. 3E-H**). While network activity in naive mice contains no stimulus specific information (**Fig. 3E**), on learned days, increases in activity of the network temporally coordinate with the preferred stimulus (**Fig. 3F**). In nonpreferred stimuli in learned days, there is also temporal coordination in network activity with the preferred stimulus, but only until approximately 0.7 s into the period, becoming significant at 0.8 s, and at which time there begins strong suppression of network activity that continues until reaching a plateau until approximately 2 s (**Fig. 3F,H**). Activity in both preferred and nonpreferred conditions in learned days suggests much higher temporal coordination of activity than of that in both conditions in naive days (**Fig. 3G,H**).

We next examined max spiking activity by cell as a function of time. We find that when bounded by the preferred stimulus period (0 to 1.4 s), there are significant differences in the active task between naive and learned preferred and nonpreferred max spikes, and learned preferred and nonpreferred max spikes (**Fig. 4A**) (two-sided Kolmogorov-Smirnov tests with Bonferroni Correction: new Alpha = .0125. Pref naive vs. NP naive: *D*(1016) = .0629, *p* = .2585. Pref learned vs NP learned: *D*(856) = .1462, *p* = 1.21×10^−4^. Pref naive vs. pref learned: *D*(938) = .1778 *p* = 3.37×10^−7^. NP naive vs. NP learned: *D*(936) = .2382, *p* = 4.36×10^−12^). These results are in accord with the largest changes in mean network activity within 0 to 1.4 s limited to those in the nonpreferred stimulus. When we expand the time window of max spiking to that of the nonpreferred stimulus (0 to 4.2 s), the difference between naive and learned sessions in the nonpreferred stimulus is magnified (**Fig. 4B**) (two-sided Kolmogorov-Smirnov test; NP naive vs. NP learned: *D*(936) = .4041, *p* = 5.42×10^−34^). By approximately 0.45 s, 50% of cells have had their max spikes, a change from the naive day in which 50% of cells had their max spikes by about 1.8 s, which is roughly the half-way point of the nonpreferred stimulus period. At 1.4 s, the period of the preferred stimulus, 70% of cells have had their max spikes in the learned day. We believe that this bias toward early activity in the nonpreferred stimulus in learned days is attributable to the network’s learning of and preference for the preferred stimulus. Activity in both conditions ramps in anticipation of the preferred stimulus; when the network identifies that the stimulus is the nonpreferred – approximately at 0.6 to 0.8 s – activity is dampened.

**Figure 4.**
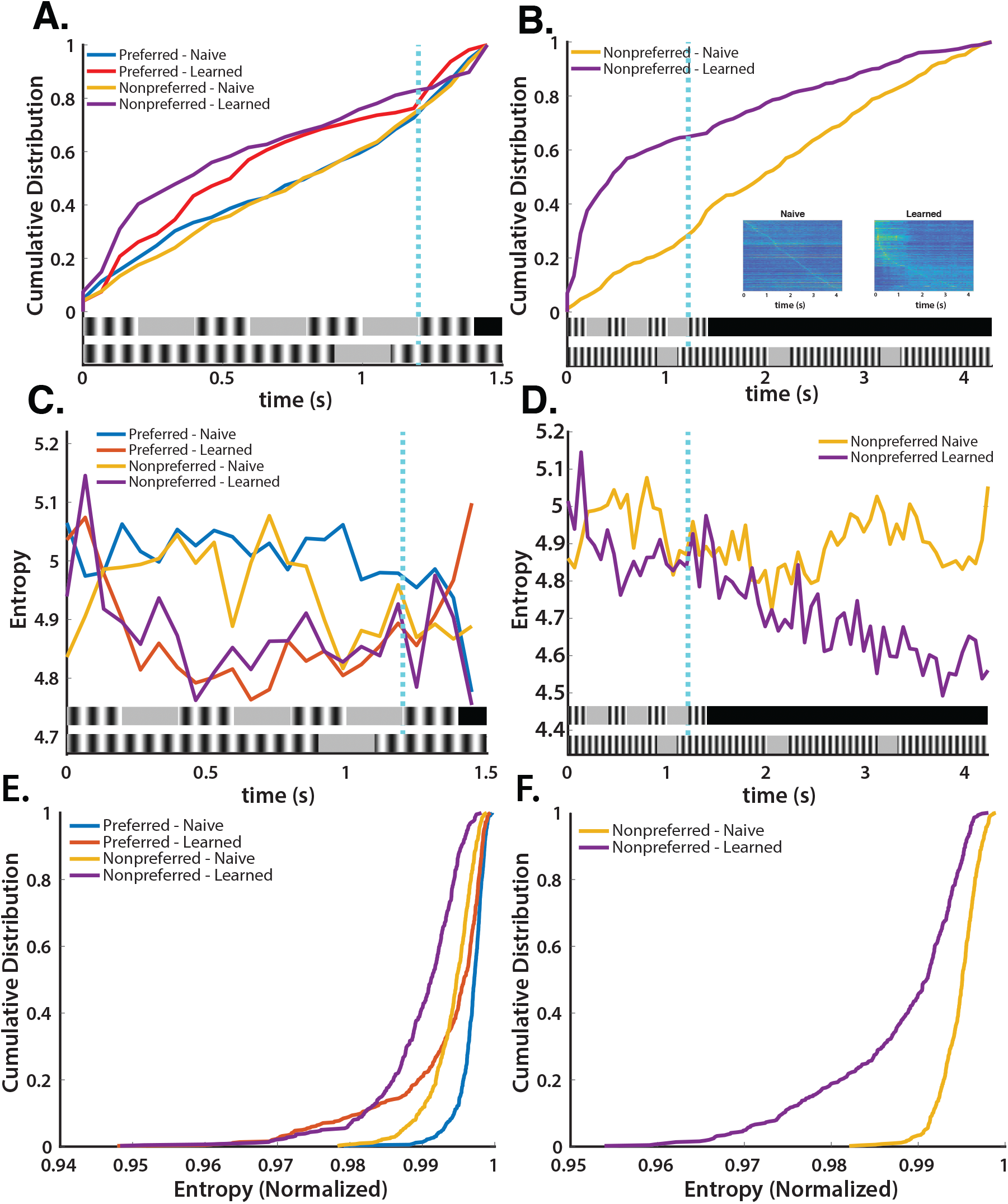
Distinct profiles of stimulus specific network activity emerge with learning. **A**. Cumulative Distribution of max spikes in the preferred stimulus period (0 to 1.4 s) shows earlier activity in learned days stimuli than naive days and significant differentiation between preferred and nonpreferred stimuli in learned days (two-sided Kolmogorov-Smirnov tests with Bonferroni Correction: new Alpha = .0125. Pref naive vs. NP naive: *D*(1016) = .0629, *p* = .2585. Pref learned vs NP learned: *D*(856) = .1462, *p* = 1.21×10^−4^. Pref naïve vs. pref learned: *D*(938) = .1778 *p* = 3.37×10^−7^. NP naive vs. NP learned: *D*(936) = .2382, *p* = 4.36×10^−12^). **B**. Cumulative Distribution of max spikes in the nonpreferred stimulus period (0 to 4.2 s) shows significantly earlier activity in learned days than in naive days (two-sided Kolmogorov-Smirnov test; NP naive vs. NP learned: *D*(936) = .4041, *p* = 5.42×10^−34^). Insets show average network activity sorted by max spikes in each day. **C**. Entropy of mean network activity as a function of time (0 to 1.4 s) shows decreased variability in learned days in both stimuli from naive days. **D**. Entropy of mean network activity as a function of time (0 to 4.2 s) shows decreased variability in learned days. **E**. Cumulative Distribution of entropy of each cell’s preferred times of firing in network decreases in learned days from naive days in the preferred stimulus period (0 to 1.4 s) (two-sided Kolmogorov-Smirnov tests with Bonferroni Correction: new Alpha = .0125. Pref naive vs. NP naive: *D*(1016) = .4656, *p* = 4.08×10^−49^. Pref learned vs NP learned: *D*(856) = .4545 *p* = 1.36×10^−39^. Pref naive vs. pref learned: *D*(936) = .3121, *p* = 1.89×10^−20^. NP naive vs. NP learned: *D*(936) = .3798, *p* = 4.42×10^−30^). **F**. Cumulative Distribution of entropy of each cell’s preferred times of firing in network decreases in learned days from naive days in the nonpreferred stimulus period (0 to 4.2 s) (two-sided Kolmogorov-Smirnov test; NP naive vs. NP learned: *D*(936) = .4898, *p* = 9.63×10^−50^).

To further characterize leaning, we calculated the entropy of total network activity as a function of time in the preferred condition (**Fig. 4C**). Sustained higher entropy values were seen in naive days in both conditions. In learned days, entropy values were initially high and then plateaued beginning at approximately 0.6 s, and then slightly increased at about 1.3 s, which follows the water reward. This demonstrates that the network’s activity was more uniform trial-to-trial in learned days than in naive days in a time-dependent manner, which both suggests learning and further implicates 0.6 to 0.8 s as the period at which discrimination of stimuli begins in learned days.

We then expanded the time window to the nonpreferred period, 0 to 4.2 s, to compare the entropy values between the naive and learned days in the nonpreferred condition (**Fig. 4D**). We find a steeper decrease in entropy over time in the learned days than in the naive days. This is again likely attributable to more uniform activity in the learned day due to sustained suppression of network activity in the nonpreferred condition. To examine emergence of uniformity of the network activity for specific periods (defined by the stimulus) we next calculated the normalized entropy of each cell’s firing time. We find that in learned days entropy across the network in both preferred and nonpreferred conditions entropy is left-shifted significantly (**Fig. 4E**) (two-sided Kolmogorov-Smirnov tests with Bonferroni Correction: new Alpha = .0125. Pref naive vs. NP naive: *D*(1016) = .4656, *p* = 4.08×10^−49^. Pref learned vs NP learned: *D*(856) = .4545 *p* = 1.36×10^−39^. Pref naive vs. pref learned: *D*(936) = .3121, *p* = 1.89×10^−20^. NP naive vs. NP learned: *D*(936) = .3798, *p* = 4.42×10^−30^). Likewise during the nonpreferred period, there is a left-shifted distribution in learned compared to naive days (**Fig. 4F**) (two-sided Kolmogorov-Smirnov test; NP naive vs. NP learned: *D*(936) = .4898, *p* = 9.63×10^−50^). The uniformity of cells’ firing times in both time windows increases in learned days suggesting learning of stimuli.

### Learning is accompanied by correlated recruitment of network activity to both stimuli

Upon establishing that there were task-dependent changes to network activity that accompany learning, we investigated to what extent network activity predicted stimulus type. We again used a bootstrapped SVM to predict stimuli from cellular activity in naive and learned days. Similar to the SVM we used to predict stimuli from licking behavior, we run a bootstrapped SVM of 1000 iterations per time bin (.067 s time bins; see Methods). We find that in naive days, there is moderate but sustained predictability from 0.3 to 1 s, which is the period in which the two stimuli diverge (**Fig. 5A**). In learned days, predictability begins to ramp at 0.7 s eventually becoming more predictable than the naive day at 1 s. These results demonstrate that network activity differs in a time-dependent manner between the two days.

**Figure 5.**
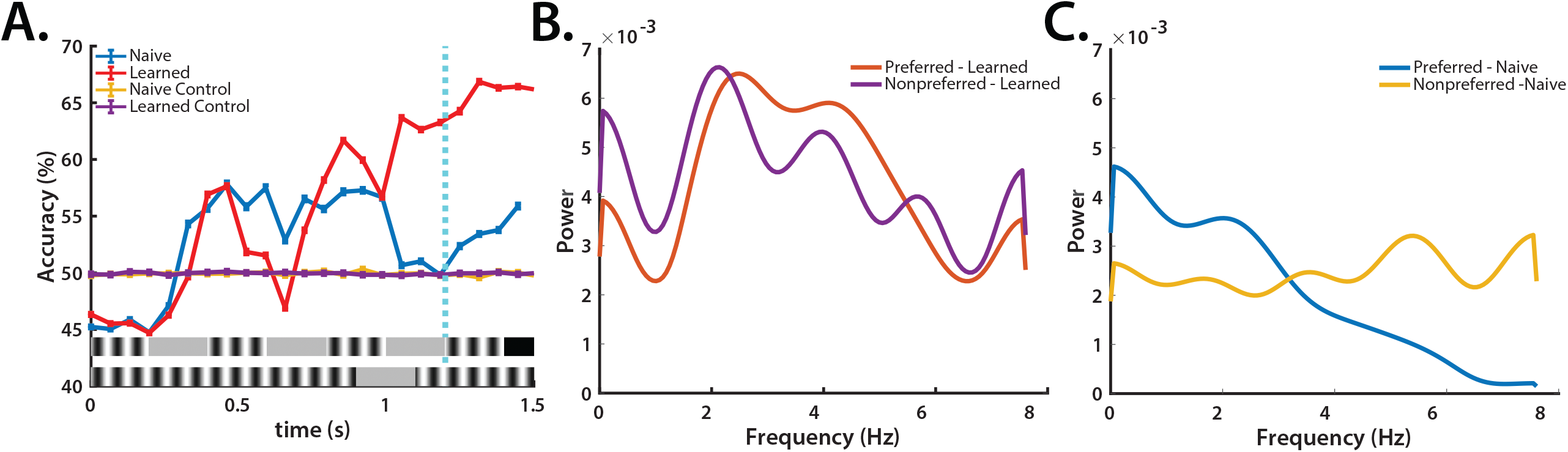
Learning is accompanied by correlated recruitment of network activity to both stimuli. **A**. Support Vector Machine (SVM) increasingly predicts stimulus from neural data in learned but not naive days. **B**. Transform of oscillatory activity in learned day shows preference for 2.5 Hz (preferred stimulus period) in preferred and nonpreferred stimulus. **C**. Transform of oscillatory activity in naive day shows moderate preference for 2.5 Hz (preferred stimulus period) in preferred stimulus, but not nonpreferred stimulus.

Based upon our earlier analyses in which we found similarities in network activity between the preferred and nonpreferred conditions in learned days, we hypothesized that the same network was recruited in both conditions and that it was tuned to the preferred stimulus. We thus interpret the higher predictability in naive days in the SVM from 0.3 to 1 s as representative of two distinct networks responding to two different stimuli. In learned days, this distinction is effaced until the decision point – approximately 0.7 s – at which point there is suppression of network activity in the nonpreferred stimulus and sustained activity in the preferred stimulus.

We tested this hypothesis by performing a power spectral density analysis of average network activity by day and by stimulus. If indeed the same network is recruited in both conditions in the learned day, power should be highest in the 2.5 Hz range (the frequency of the preferred stimulus) prior to the decision point. In naive days, the highest powers should differ between stimuli as the networks are responding differentially. We find that indeed, there is high correlation between power spectra prior to the decision point in both conditions in learned days with the highest power centered at approximately 2.5 Hz (**Fig. 5B**). Conversely, power spectra are not correlated in naive days and only in the preferred stimulus is power highest at approximately 2.5 Hz (**Fig. 5C**). Thus, we conclude that learning is accompanied by changes in network activity that anticipates the preferred stimulus regardless of whether the preferred or nonpreferred stimulus is presented, and only at the decision point does network activity diverge.

## Discussion

Using a go/no-go audio-visual timing task, we have demonstrated that mice learned to perform the TPSD task successfully as assessed through an increase in discriminability index and tuning of licking profiles. Learned performance was attributable to changes in response to the nonpreferred temporal pattern in which licking was suppressed early into the stimulus period. Similar results were seen in neural activity in V1, L-2/3 in which activity was suppressed in the nonpreferred stimulus but was released in a temporally defined manner in the preferred stimulus. Interestingly, learning was associated with the emergence of neural resonance of 2.5 Hz that reflected the temporal structure of the preferred stimulus. The robust neural resonance was associated with neural trajectories evoked by both preferred and non-preferred stimuli. This shared oscillatory activity contributed to the reduced accuracy of the decoder early in the stimulus. The eventual increase in decoder accuracy likely results from the refinement of emergent neural activity in V1 to the temporal structure of stimuli. Thus, our data shows that temporal rhythms are encoded by a combination of emergent neural oscillatory activity followed by refinement of neural dynamics to reflect stimulus specific information – thus supporting the intrinsic model of timing in which time is encoded by network dynamics (**Fig. S1**).

Due to the complexity of understanding temporal processing, time has been categorized into distinct types such as sensory versus motor timing and interval versus pattern timing, as well as distinguishing between different time-scale ranges (Paton and Buonomano, 2018). For example, many tasks in interval timing require the identification of isolated segments of time such as waiting to cross a street or identifying a single musical note. While interval timing can be studied as its own entity, it is also important in pattern timing. Pattern timing is composed of intervals and contains a temporal structure (Paton and Buonomano, 2018). For instance, to understand language, one must recognize the overall prosody of speech, as well as the pauses between words. The time scale in which interval and pattern timing occurs is on the order of tens of milliseconds to a few seconds, although it is unknown if the neural mechanisms of interval and rhythmic timing are shared and if intrinsic timing mechanisms that contribute to interval timing also contribute to learning of temporal patterns (Hardy and Buonomano, 2016).

Although early theories of timing proposed centralized mechanisms dedicated entirely to temporal perception, it has since been established that different neural mechanisms are involved in processing time at different time-scales (Paton and Buonomano, 2018). However, it has yet to be determined if the mechanisms of temporal perception in the range of seconds and subseconds are distributed across brain regions or if local networks within different regions can intrinsically encode time, albeit through a diversity of network dynamics (Zhou et al., 2020; Zhou et al., 2022). While the locus of sub second and second timing and temporal predictions has been attributed to higher order cortical areas (Leon and Shadlen, 2003; Jazayeri and Shadlen, 2015; Licata et al., 2017) and subcortical areas ((Bakhurin et al., 2017; Zhou et al., 2020; Toso et al., 2021), accumulating evidence suggests that primary visual cortex (V1) exhibits response modulation to “higher” functions such as spatiotemporal learning, reward prediction, and attention (Gavornik and Bear, 2014b). A few studies have examined the role of visual cortical neurons in timing (Duysens et al., 1996; Shuler and Bear, 2006; Namboodiri et al., 2015). One study in rats showed that as rats perform a visually cued timing task, the V1 cortical activity rapidly modulated to predict arrival of reward (Shuler and Bear, 2006) and cholinergic function contributed to the modification of V1 activity (Chubykin et al., 2013). Another study using a similar task showed that indeed cortical activity in V1 reflected the duration of a target interval (Namboodiri et al., 2015). Further, responses in V1 can be modulated to predict a stimulus at a spatial location (Fiser et al., 2016). One interpretation of our data is that the evolution of neural resonance early on in the patterned stimuli predicts arrival of reward, suggestive of a novel mechanistic driver of temporal predictions in V1.

Our future studies are aimed at understanding the network mechanisms that drive refinement of neural activity that allows discrimination of patterns of stimuli. Complex interplay of synaptic excitation and inhibition allows cortical neurons to adaptively respond to sensory stimuli, discriminate between stimuli, and integrate sensory inputs (Isaacson and Scanziani, 2011; Ferguson and Cardin, 2020). Converging evidence across many studies and model systems, including our own work, shows that selectivity to the interval of a stimulus duration is the result of dynamic shifts in excitation-inhibition (E-I) balance (Edwards et al., 2007; Kostarakos and Hedwig, 2012; Goel and Buonomano, 2016). Consistent with previous *in-vitro* work in cortical slices (Goel and Buonomano, 2016), our data suggests that network suppression is one potential mechanism that drives learning. Encoding of intervals and patterns in sensory cortical circuits can result from changes in the E-I ratio at temporally defined periods (Goel and Buonomano, 2016) and by multiple interneuron populations (Cardin, 2018). V1, L-2/3 is composed primarily of Pyramidal (Pyr) cells which are synapsed by PV cells at the cell body and axonal hillock (Gonchar and Burkhalter, 1997). Somatostatin (SST) cells synapse onto pyramidal cells’ dendrites thus providing more fine-tuned inhibition. Both PV and SST cells have been implicated in short term plasticity, which is one of the mechanisms proposed to drive sub second sensory timing (Buonomano, 2000; Motanis et al., 2018; Seay et al., 2020). PV cells provide reliable inhibition within a short window which can help constrain pyramidal cell firing to temporally defined windows (Pouille and Scanziani, 2001; Cardin et al., 2010). We speculate that inhibition, driven by PV cell activity, which is broadly tuned due to their anatomical connectivity, can drive temporal encoding of rhythmic patterns. Indeed, in a recent study, PV neurons were shown to be important in mediating reward timing (Monk et al., 2020). However, SST cells also modulate cortical output. SST neurons not only provide dendritic pyramidal cell inhibition but also control PV cell output (Atallah et al., 2012). Therefore a complex interaction between SST and PV cells determines the balance between somatic and dendritic inhibition on pyramidal cells (Xue et al., 2014), thus contributing to temporal encoding (Cardin, 2018). In addition, VIP interneurons inhibit PV and SST interneurons, thereby reducing tonic inhibition of target pyramidal neurons. Through this disinhibitory circuit, VIP cells can increase the gain of pyramidal neurons particularly during reinforcement learning (Letzkus et al., 2011; Jiang et al., 2013; Pi et al., 2013). Therefore in our goal directed TPSD task, VIP cells can restrict PV and SST function thus influencing learning. Further, VIP cells receive functional input from anterior cingulate cortex (ACC), which is implicated in attentional modulation of sensory discrimination. One study showed that ACC input contributes to spatially driven predictive responses of V1 (Fiser et al., 2016). Thus, VIP cells could be effective drivers of temporal encoding in a goal driven task. However, we also want to emphasize that circuits in V1 are not the only contributors to TPSD task and that multiple areas such as auditory cortex, ACC, and other downstream areas are likely involved.

Our results thus far suggest that the emergence of complex neural dynamics in V1 accompanies temporal pattern learning. This opens the door to future studies probing the mechanistic substrates of excitation and inhibition that fine tune temporal learning, and offers insights into translational studies of time perception. Indeed, an important hallmark of learning is being sensitive to and remembering the temporal structure of events so that we can make predictions and guide our future decisions. As a result, it is not surprising that disruptions in timing and timed performance are associated with a number of neurological disorders such as Parkinson’s, Schizophrenia, and Autism Spectrum Disorder. Thus, by combining cutting-edge tools with simple behavior we not only provide fundamental information about neural mechanisms of timing but provide an assay to test deficits in timing.

## Supplementary Figure Legends

**Figure S1.**
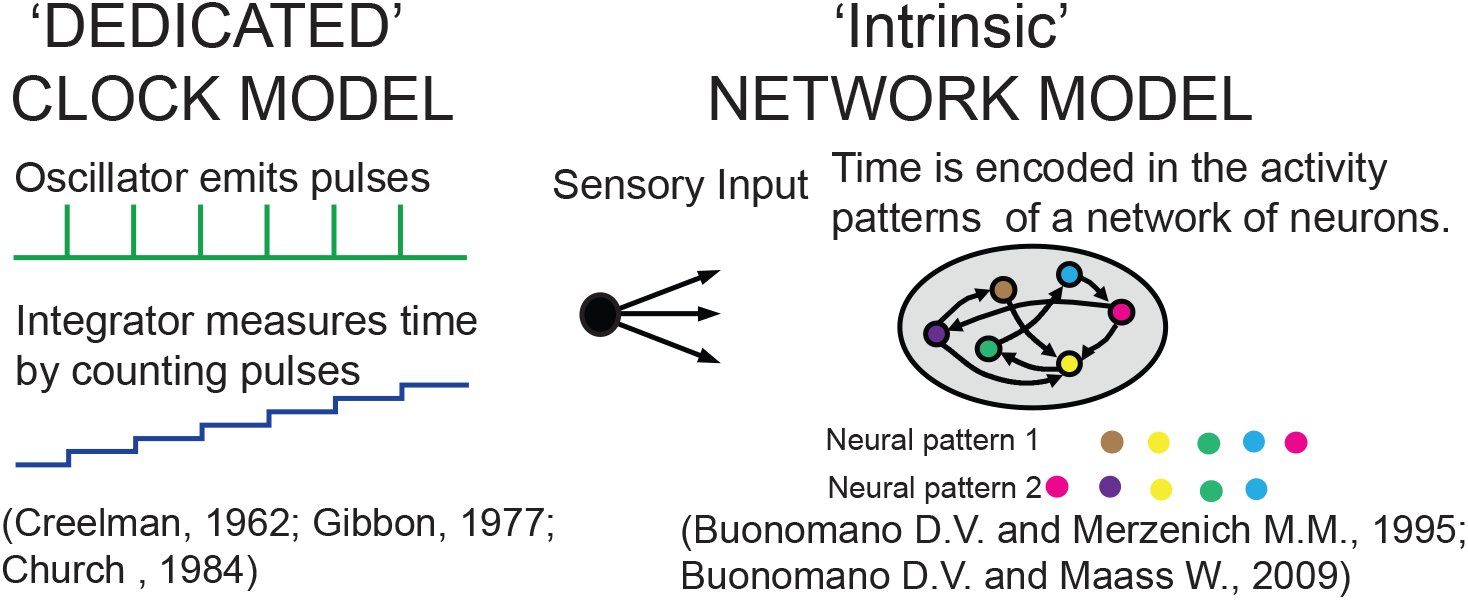
Models of neural mechanisms of timing. Dedicated models propose centralized mechanisms in which an oscillator generates periodic pulses which are then counted by a downstream integrator. Intrinsic models propose unique network dynamics as the means by which time is encoded.

**Figure S2.**
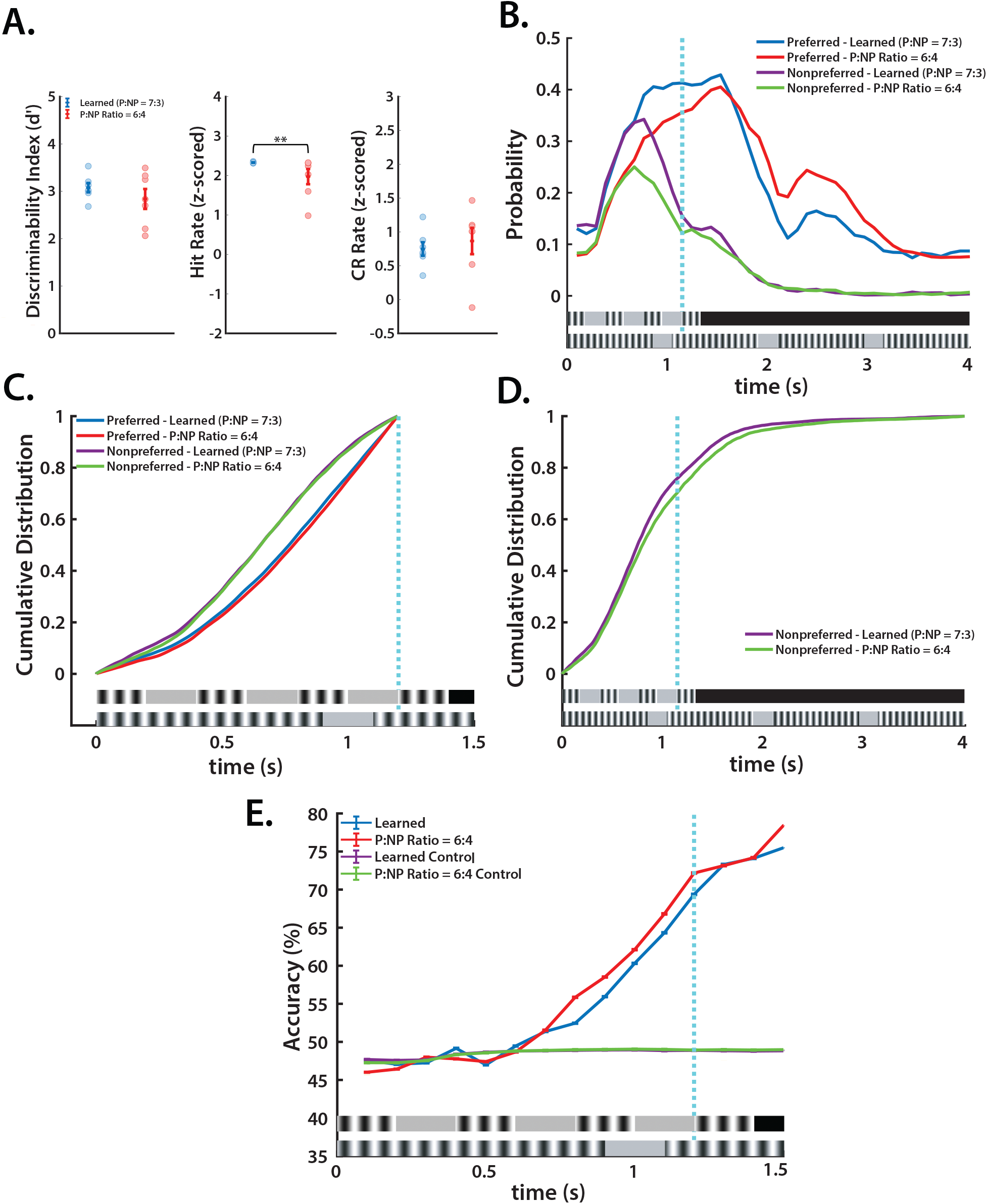
Learned performance is not dependent on the ratio of preferred: non-preferred stimuli. **A**. Performance remains similar between learned sessions (P:NP ratio = 7:3) and sessions with a 6:4 preferred: nonpreferred stimuli ratio (*Discriminability Index (d’)*: two-tailed t-test, *t*(12) = 1.03, *p* = .32. *Hit Rate*: Wilcoxon rank sum, *p* = .0093. *CR Rate*: Wilcoxon rank sum, *p* = .3829). **B**. Licking probabilities remain similar between learned sessions and sessions with a 6:4 preferred: nonpreferred stimuli ratio. **C**. CDF of licking from 0 to 1.2 s shows similarities between learned and 6:4 preferred: nonpreferred stimuli ratio sessions (two-sided Kolmogorov-Smirnov tests with Bonferroni Correction: new Alpha = .001. Pref learned vs. NP learned: *D*(86883) = .1376, *p* = 3.1×10^−276^. Pref learned vs. Pref P:NP ratio = 6:4: *D(*109563) = .0253, *p* = 3.6×10^−15^. Pref P:NP ratio = 6:4 vs. NP P:NP ratio = 6:4: *D*(65683) = .1551, *p* = 2.58×10^−294^. NP learned vs. NP Pref P:NP ratio = 6:4: *D*(43003) = .0183, *p* = .0015). **D**. CDF of licking from 0 to 4.2 s shows similarities between learned and 6:4 preferred: nonpreferred stimuli ratio sessions, but not control sessions (two-sided Kolmogorov-Smirnov test, *D*(58927) = .058, *p* = .1.58×10^−43^). **E**. Support Vector Machine (SVM) increasingly predicts stimulus type from licking in learned and P:NP ratio = 6:4 sessions.

**Figure S3.**
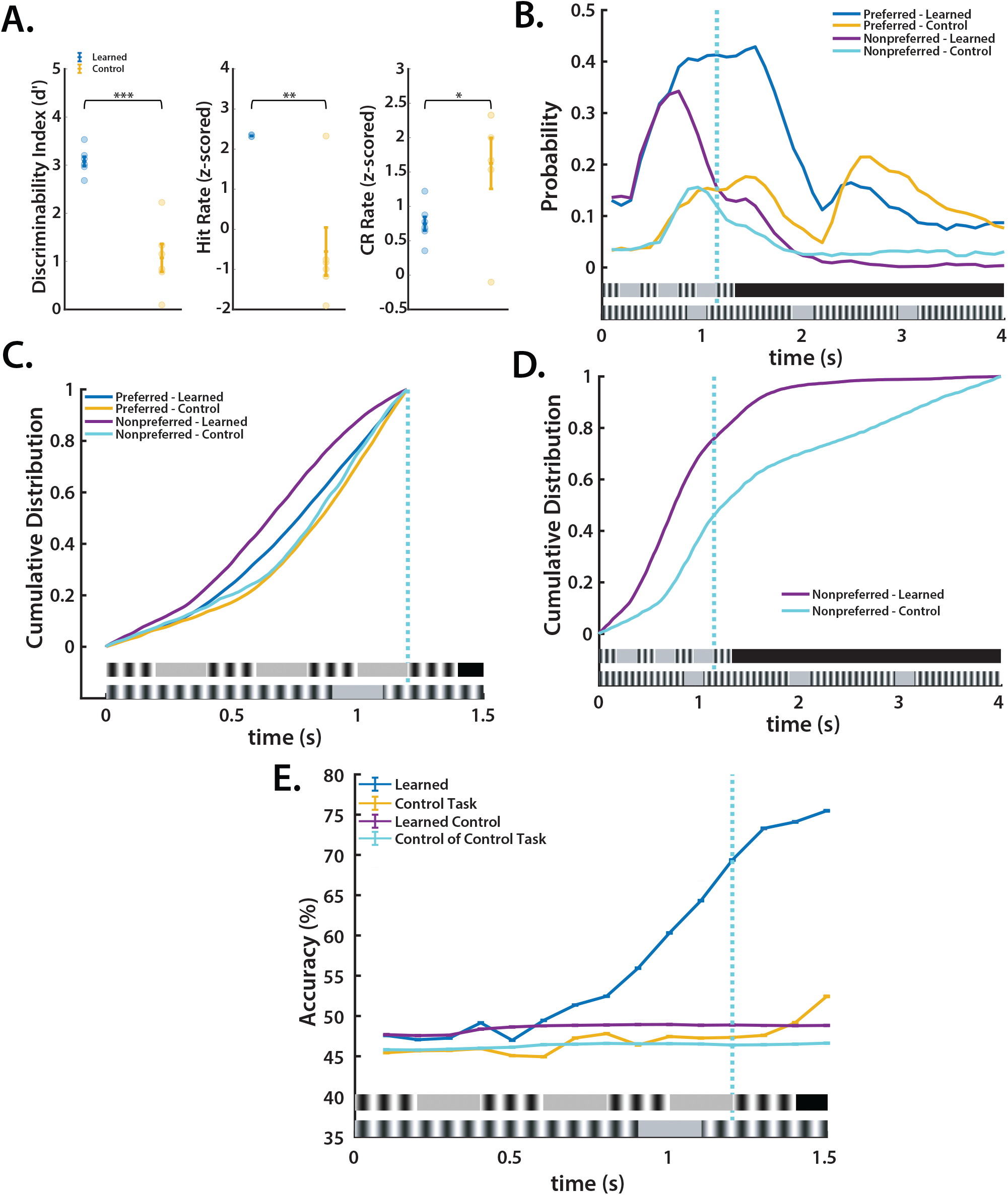
Lack of learned performance on the TPSD task in the absence of stimulus. **A**. Performance decreases between learned sessions and control sessions (*Discriminability Index (d’)*: two-tailed t-test, *t*(11) = 7.02, *p* = 2.21×10^−5^. *Hit Rate*: Wilcoxon rank sum, *p* = .007. *CR Rate*: two-tailed t-test, *t*(11) = 2.46, *p* = .03). **B**. Licking probabilities differ between learned and control sessions. **C**. CDF of licking from 0 to 1.2 s shows changes in licking between learned and control sessions (two-sided Kolmogorov-Smirnov tests with Bonferroni Correction: new Alpha = .001. Pref learned vs. NP learned: *D*(86883) = .1376, *p* = 3.1×10^−276^. Pref learned vs pref control: *D*(81272) = .1069, *p* = 2.10×10^−134^. Pref control vs NP control: *D*(23039) = .0435, *p* = 9.91×10^−8^. NP learned vs NP control: *D*(28650) = .2245, *p* = 1.25×10^−208^). **D**. CDF of licking from 0 to 4.2 s shows changes in licking between learned and control sessions (two-sided Kolmogorov-Smirnov test, *D*(42991) = .3259, *p* → 0). **E**. Support Vector Machine (SVM) increasingly predicts stimulus type from licking in learned sessions, but not remains at chance through control sessions.

**Figure S4.**
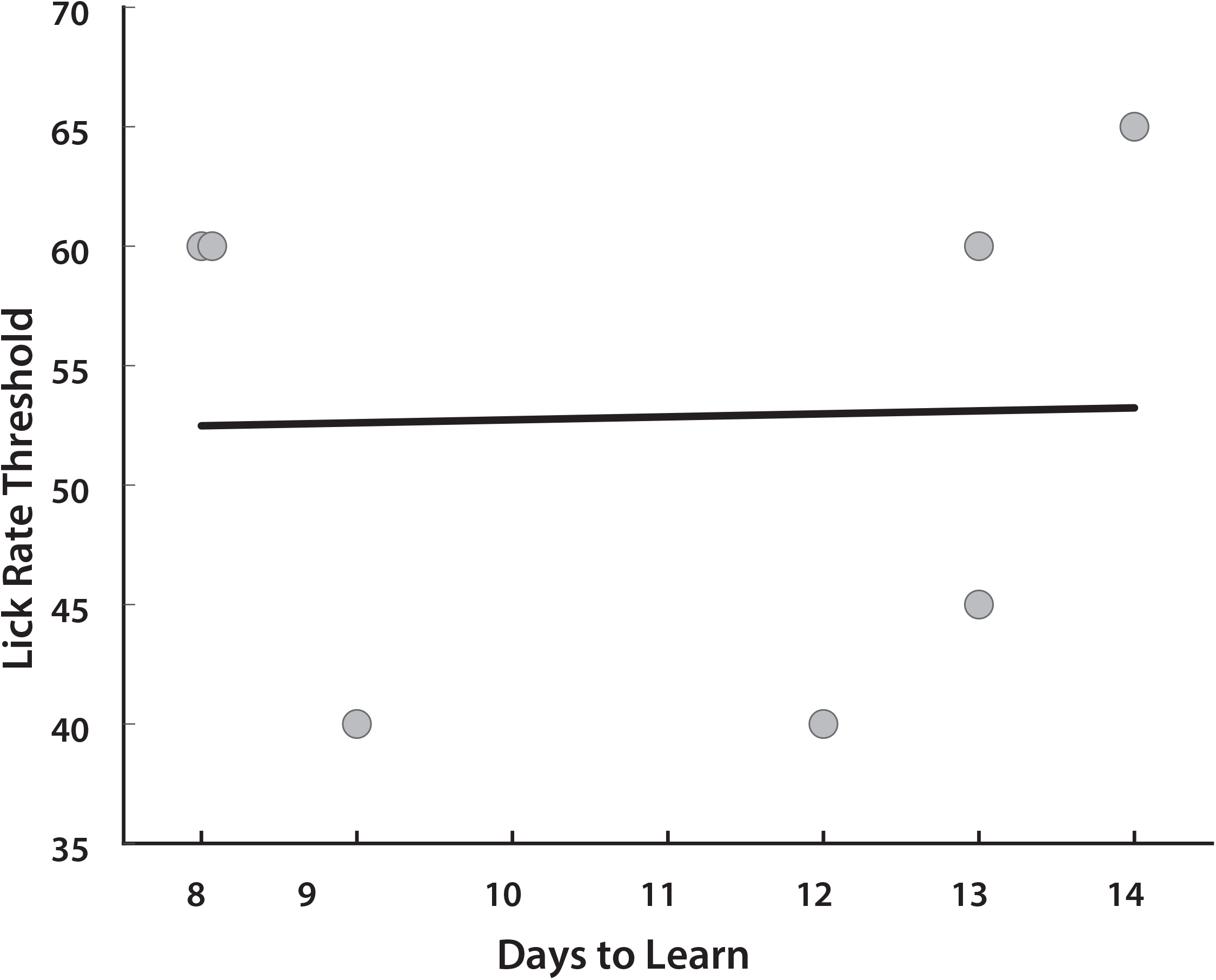
Individualized lick rate thresholds do not affect time to learning (Pearson’s r, *r*(12) = .03, *p* = .949).

## Author contributions

S.P. and A.G. conceived the project and designed the experiments. S.P. wrote the MATLAB code for data acquisition and analysis with help from A.G. S.P, W.M., O. AW conducted the experiments and S.P analyzed the data. A.G. and S.P interpreted the data and wrote the paper with input from other authors.

## Acknowledgements

The authors thank Aaron Seitz, Ladan Shams and Dean Buonomano for discussions that helped improve the paradigm; Edward Zagha for technical expertise with Power Spectrum density Analysis.

## Materials and Methods Experimental Animals

All experiments followed the U.S. National Institutes of Health guidelines for animal research, under animal use protocols approved by the Chancellor’s Animal Research Committee and Office for Animal Research Oversight at the University of California, Riverside (ARC #2019-0036). We used male and female FVB.129P2 WT mice (JAX line 004828). All mice were housed in a vivarium with a 12/12 h light/dark cycle and experiments were performed during the light cycle. The FVB background was chosen because of its robust breeding.

### Go/No-go temporal pattern sensory discrimination (TPSD) task for head restrained mice

Awake, head-restrained young adult mice (2-4 months) were allowed to run on an air-suspended polystyrene ball while performing the task in our custom built rig (**Fig. 1A**). Prior to performing the task, the animals were subjected to handling, habituation, and pretrial phases. After recovery from headbar/cranial window surgery, mice were handled gently for 5 min every day, until they were comfortable with the experimenter and would willingly transfer from one hand to the other to eat sunflower seeds. This was followed by water deprivation (giving mice a rationed supply of water once per day) and habituation to the behavior rig. During habituation, mice were head-restrained and acclimated to the enclosed sound-proof chamber and allowed to run freely on the 8 cm polystyrene ball. Eventually, mice were introduced to the lickport that dispensed water (3-4 μL) and recorded licking (custom-built at the UCLA electronics shop), followed by the audio-visual stimuli. This was repeated for 10 min per session for 3 days. Starting water deprivation prior to pretrials motivated the mice to lick (Guo et al., 2014). After habituation and ∼15% weight loss, mice started the pretrial phase of the training. During pretrials, mice were shown the preferred stimulus only with no punishment time associated with incorrect responses. This was done in order to 1) teach the mice the task structure 2) encourage the mice to lick and to remain motivated. The first day consisted of 150 trials and subsequent days of 250. The reward, as in the TPSD main task, was dispensed at 1.2 s and remained available to the mice until 2 s, at which time it was sucked away by a vacuum. The mice were required to learn to associate a water reward soon after the stimulus was presented and that there was no water reward in the inter-trial interval (4 s period between trials). Initially, during pre-trials the experimenter pipetted small drops of water onto to the lickport to coax the mice to lick. Once the mice learned this and licked with 85% efficiency, they were advanced to the go/no-go task.

The TPSD task is a go/no-go task composed of two sequences of synchronous audio-visual stimuli (**Fig. 1B**). Visual stimuli are 90° drifting sinusoidal gratings and are accompanied by a synchronous 5 kHz tone at 80 dB. Within each sequence, four stimuli are presented that differ only in temporality. Our preferred sequence is composed of 4 stimuli of 200 ms; our nonpreferred sequence is composed of 4 stimuli of 900 ms. Each set of the sequences is separated by a 200 ms period of silence accompanied by a grey screen. A water reward is dispensed at 1.2 s and remains available until 2 s, at which time it is sucked away by a vacuum. A custom built lickport (UCLA engineering) dispensed water, vacuumed it, and recorded licking via breaks in an infrared (IR) beam. Breaks were recorded at 250 Hz. The window in which mice’s licking count toward a response is 1 to 2 s in both stimuli. A time out period (6.5 to 8 s), in which the monitor shows a black screen and there is silence, is instituted if the mouse incorrectly responds. The first session was composed of 250 trials, and subsequent days of 350. Depending on the stimulus presented, the animal’s behavioral response was characterized as “Hit”, “Miss”, “Correct Rejection” (CR) or “False Alarm” (FA) (**Fig. 1B**). An incorrect response resulted in the time-out period .

To expedite learning, we set the ratio of preferred to nonpreferred stimuli to 70:30 as we found that mice are more prone to licking (providing a ‘yes’ response) than to inhibiting licking (providing a ‘no’ response). We additionally instituted an individualized lick rate threshold to encourage learning as we found that lick rates differed significantly from mouse to mouse. **Fig. S4** shows the licking thresholds (calculated from lick rates) for mice and shows no significant correlation between licking thresholds and learning rates. This indicates that the individualized lick rate threshold was used as a learning aid to complete the task and did not affect their learning rates or their reliance on the stimulus for task completion. To confirm that mice learned rather than took advantage of the biased 70:30 preferred to nonpreferred trial ratio, we tested mice for 2 additional sessions using a 60:40 ratio of preferred to nonpreferred stimuli (**Fig. S2 A-E**). We retain a greater number of preferred stimuli as the total time mice encounter preferred stimuli is less than that of encountering nonpreferred stimuli within a 60:40 trial session (294 s vs 588 s respectively). Following, mice performed a control task, during which visual and auditory stimuli were not presented. Our data shows that mice did not show a leaned performance, indicating that they relied on the sensory stimuli for task completion (**Fig. S2 A-E**).

Custom-written routines and Psychtoolbox in MATLAB were used to present the visual stimuli, to trigger the lickport to dispense and retract water, and to acquire data.

### Cranial window surgery

Craniotomies were performed at 6-8 weeks as previously described(Mostany and Portera-Cailliau, 2008; Goel et al., 2018). Mice were given dexamethasone (0.2 mg/kg) and carprofen (5 mg/kg) intraperitoneally and subcutaneously respectively. Mice were anesthetized with isoflurane (5% induction, 1.5-2% maintenance via nose cone) and placed in a stereotaxic frame. Under sterile conditions, a 4.5 mm diameter craniotomy was drilled over the right primary visual cortex (V1) and covered with a 5 mm glass coverslip. Before securing the cranial window with a coverslip, we injected ∼60-100 nl of pGP-AAV-syn-jGCaMP7f-WPRE. A custom U-shaped aluminum bar was attached to the skull with dental cement to head restrain the animal during behavior and calcium imaging. For two days following surgery, mice were given dexamethasone (0.2 mg/kg) daily.

### Viral constructs

pGP-AAV-syn-jGCaMP7f-WPRE were purchased from Addgene and diluted to a working titer of 2e^13^ with 1% filtered Fast Green FCF dye (Fisher Scientific).

### In-vivo two photon calcium imaging

Calcium imaging was performed on a Scientifica 2-photon microscope equipped with a Chameleon Ultra II Ti:sapphire laser (Coherent), resonant scanning mirrors (Cambridge Technologies), a 20X objective (1.05 NA, Olympus), multialkali photmultiplier tubes (R3896, Hamamatsu) and ScanImage software (Pologruto et al., 2003). Prior to calcium imaging, head-restrained mice were habituated to a sound-proof chamber and allowed to run freely on a polystyrene ball (**Fig. 1A, 3A**). Visually evoked responses of L2/3 pyramidal cells from V1 were recorded at 15 Hz in 1 field of view. Each field of view (FOV) consisted of a mean of 117.25 pyramidal cells (sd = 34.01). In each animal, imaging was performed at 150-250 μm.

### Data analysis

#### Discriminability index (d’) and CR and Hit Rates

d’ was calculated using the MATLAB function *norminv*, which returns the inverse of the normal cumulative distribution function:

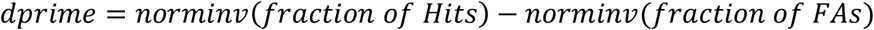

If either rate reached 100% or 0%, we arbitrarily changed the value to either 99% or 1%, respectively. We did this to avoid generating z-scores of infinity that would inaccurately characterize the mice’s performance

#### Licking Probabilities

Probabilities were taken by binning licks per 0.1 s window per trial per mouse. We then averaged each mouse’s probability per time to generate a distribution of probabilities based on trial session and stimulus type. We use the best 150 trials from each day and each mouse as determined by the discriminability index (*d’*).

#### Data analysis for calcium imaging

Calcium-imaging data were analyzed using suite2p (Pachitariu M et al., 2017) and custom-written MATLAB routines. All data was then processed using suite2p for image registration, ROI detection, cell labeling, and calcium signal extraction with neuropil correction. Once Suite2P had performed a rigid and non-rigid registration and then detected regions-of-interest (ROIs) using a classifier, we manually selected cells using visual inspection of ROIs and fluorescence traces to ensure the cells were healthy. We then used the deconvolved spikes determined by suite2p in our subsequent analysis that used custom-written MATLAB scripts.

#### Network and Cellular Entropy

Network entropy was calculated using Shannon’s entropy equation, in which *H(X)* represents the entropy of network activity per time bin; *n* represents the total number of network activity values; *i* represents a given element in *n*; and *x*_*i*_ represents the network activity value of the *i*th element:

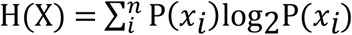

Network activity per time bin per trial was averaged on a 0 to 1 scale and then rounded to two decimal places so as to calculate approximate but meaningful entropy values (e.g. if network activity was highly uniform in the hundredths digit but not in the thousandths, an entropy value may have been as high as a time bin in which network activity was uniform in the tenths but not the hundredths digit). Then an entropy value for each time bin was calculated.

Cellular entropy was calculated using Shannon’s entropy equation with a normalization. A normalization was performed to account for differing firing rates among cells. *H(X)* represents the entropy of a given cell’s time preference for firing within a set window. Two windows were used: 0 to 1.4 s and 0 to 4 .2 s, the periods of the preferred and nonpreferred stimuli respectively. *n* represents the total number of time periods in which a cell fired; *i* represents a given element in *n*; and *x*_*i*_ represents the time period of the *i*th element. We normalize using the maximum entropy value, *log*_*2*_n.

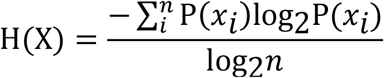

#### Power Spectral Density of Network Activity

Power spectra of network activity were calculated by taking the mean activity of the network over a given time. We then used the *pmtm* function in MATLAB to derive power spectra.

#### Support Vector Machine (SVM)

We used the SVM available in the MATLAB Machine Learning and Deep Learning toolbox via the function *fitcsvm*. We used a radial basis function as the kernel. 80% of our data was applied to training the machine and 20% applied to testing it. Instead of training one machine, we developed a strategy wherein we performed a bootstrapped SVM per time per mouse. This allowed us to generate a distribution of accuracy percentages per time such that we could locate critical times of difference during stimulus presentation. 10000 machines were generated per time per mouse for the licking predictor and then averaged as one grand distribution. 1000 machines were generated per time per mouse for the imaging predictor then averaged as one grand distribution. The fewer number of machines for the imaging predictor was due to computational constraints. The licking predictor consisted of binning licks per 0.1 s window per trial per mouse with the stimulus type as the outcome. The imaging predictor was the network’s activity with the stimulus type as the outcome in 0.067 s time bins. For a given mouse, each cell’s putative spiking activity was used as a feature space per a given time. With our licking data, we performed no pretraining optimization as we essentially were testing the accuracy of individual features, i.e. time bins of licking. With our neural data, we likewise did not perform a feature selection procedure as we found it too computationally intensive. Thus, each cell’s activity contributes to the fitting of the hyperplane. However, we performed a cross validation with 5 k-folds and used the default hyperparameter optimization algorithm in MATLAB prior to training.

### Statistical analyses

Statistical analysis of normality (Lilliefors) were performed on each data set and depending on whether the data significantly deviated from normality (p<0.05) or did not deviate from normality (p>0.05) appropriate non-parametric or parametric tests were performed. The statistical tests performed are mentioned in the text and the legends. For parametric two-group analyses, a Student t-test (paired or unpaired) was used; for parametric multi-group analyses, a one-way ANOVA was used. For non-parametric tests, we used the following: Wilcoxon rank sum test (two groups), Kolmogorov-Smirnov test (two groups), and Kruskal-Wallis test (multi-group). When multiple two group tests were performed, a Bonferroni Correction was applied to readjust the Alpha value. In the figures, significance levels are represented with the following convention: * for p < Alpha; ** for p < Alpha/10, *** for p < Alpha/100. Alpha values are .05 unless otherwise specified. Effect sizes were computed using either Cohen’s D or Cliff’s delta following Lilliefors tests of normality. In all the figures, we plot the standard error of the mean (s.e.m.). Graphs either show individual data points from each animal or group means (averaged over different mice) superimposed on individual data points.

### Exclusion of mice

5 WT mice were excluded from the data because the mice lost > 25% body weight (a criterion we established a priori). This had adverse effects on their health that was manifested in listlessness, reduced grooming and interaction with cage mates, and occasionally, seizures.

## Data availability

All the analyzed data reported in this study is available from the corresponding author upon request.

## Code availability

All code used in this manuscript is available from the corresponding author upon request.

